# Impulse responses reveal unimodal and bimodal access to visual and auditory working memory

**DOI:** 10.1101/623835

**Authors:** M. J. Wolff, G. Kandemir, M. G. Stokes, E. G. Akyürek

## Abstract

It is unclear to what extent sensory processing areas are involved in the maintenance of sensory information in working memory (WM). Previous studies have thus far relied on finding neural activity in the corresponding sensory cortices, neglecting potential activity-silent mechanisms such as connectivity-dependent encoding. It has recently been found that visual stimulation during visual WM maintenance reveals WM-dependent changes through a bottom-up neural response. Here, we test whether this impulse response is uniquely visual and sensory-specific. Human participants (both sexes) completed visual and auditory WM tasks while electroencephalography was recorded. During the maintenance period, the WM network was perturbed serially with fixed and task-neutral auditory and visual stimuli. We show that a neutral auditory impulse-stimulus presented during the maintenance of a pure tone resulted in a WM-dependent neural response, providing evidence for the auditory counterpart to the visual WM findings reported previously. Interestingly, visual stimulation also resulted in an auditory WM-dependent impulse response, implicating the visual cortex in the maintenance of auditory information, either directly, or indirectly as a pathway to the neural auditory WM representations elsewhere. In contrast, during visual WM maintenance only the impulse response to visual stimulation was content-specific, suggesting that visual information is maintained in a sensory-specific neural network, separated from auditory processing areas.

**Significance Statement:** Working memory is a crucial component of intelligent, adaptive behaviour. Our understanding of the neural mechanisms that support it has recently shifted: rather than being dependent on an unbroken chain of neural activity, working memory may rely on transient changes in neuronal connectivity, which can be maintained efficiently in activity-silent brain states. Previous work using a visual impulse stimulus to perturb the memory network has implicated such silent states in the retention of line orientations in visual working memory. Here, we show that auditory working memory similarly retains auditory information. We also observed a sensory-specific impulse response in visual working memory, while auditory memory responded bi-modally to both visual and auditory impulses, possibly reflecting visual dominance of working memory.

## Introduction

Working memory (WM) is necessary to maintain information without sensory input, which is vital to adaptive behaviour. In spite of its important role, it is not yet fully clear how WM content is represented in the brain, or whether sensory information is maintained within a sensory-specific neural network. Previous research has relied on testing whether sensory cortices exhibit content-specific neural activity during maintenance. While this has indeed been shown for visual memories in occipital areas (e.g., Harrison & Tong, 2009) and, more recently, for auditory memories in the auditory cortex (Huang, Matysiak, Heil, König, & Brosch, 2016; Kumar et al., 2016; Uluç, Schmidt, Wu, & Blankenburg, 2018), WM-specific activity in the sensory cortex is not always present (Bettencourt & Xu, 2016), fuelling an ongoing debate over whether sensory cortices are necessary for WM maintenance (Scimeca, Kiyonaga, & D’Esposito, 2018; Xu, 2017). However, the neural WM network may not be solely based on measurable neural activity, and it has been proposed that information in WM may be maintained in an “activity-silent” network (Stokes, 2015) - for example, changes in short-term connectivity (Mongillo, Barak, & Tsodyks, 2008). Potentially silent WM states should also be taken into account to better investigate the sensory-specificity account of WM.

Silent network theories predict that its neural “impulse” response to external stimulation can be used to infer its current state (Buonomano & Maass, 2009; Stokes, 2015). This has been shown in visual WM experiments, in which the evoked neural response from a fixed, neutral and task-irrelevant visual stimulus presented during the maintenance period of a visual WM task contained information about the contents of visual WM (Wolff, Ding, Myers, & Stokes, 2015; Wolff, Jochim, Akyürek, & Stokes, 2017). This not only suggests that otherwise hidden processes can be illuminated, but also implicates the involvement of the visual cortex in the maintenance of visual information, even when no ongoing activity can be detected. It has been suggested that this WM-dependent response profile might be not merely a byproduct of connectivity-dependent WM, but a fundamental mechanism that affords efficient and automatic readout of WM content through external stimulation (Myers et al., 2015).

It remains an open question, however, whether information from other modalities in WM is similarly organized. If auditory WM depends on content-specific connectivity changes that include the sensory cortex, we would expect a network-specific neural response to external auditory stimulation. Furthermore, it may be hypothesized that sensory information need not necessarily be maintained in a network that is detached from other sensory processing areas. Direct connectivity (Eckert et al., 2008) and interplay (Iurilli et al., 2012; Martuzzi et al., 2007) between the auditory and visual cortices, or areas where information from different modalities converges, such as the parietal and pre-frontal cortices (Driver & Spence, 1998; Stokes et al., 2013), raise the possibility that WM could exploit these connections even during maintenance of unimodal information. Content-specific impulse responses might be observed not only during sensory-specific but also sensory non-specific stimulation.

In the present study, we tested whether WM-dependent impulse responses can be observed in visual and auditory WM, and whether that response is sensory specific. We measured electroencephalography (EEG) while participants performed visual and auditory WM tasks. We show that the evoked neural response of an auditory impulse stimulus reflects relevant auditory information maintained in WM. Visual perturbation also resulted in an auditory WM-dependent neural response, implicating both the auditory and visual cortices in auditory WM. By contrast, visual WM content could only be decoded after visual, but not auditory perturbation, suggesting that visual information is maintained in a sensory-specific visual WM network with no evidence of WM-related connectivity with the auditory cortex.

## Materials and Methods

### Participants

Thirty healthy adults (12 female, mean age 21 years, range 18-31 years) were included in the main analyses of the auditory WM experiment and 28 healthy adults (11 female, mean age 21 years, range 19-31 years) of the visual WM experiment. Three additional participants in the auditory WM experiment and 8 additional participants in the visual WM experiment were excluded during pre-processing due to excessive eye movements (more than 30% of trials contaminated). Participants received either course credits or monetary compensation (8€ an hour) for participation and gave written informed consent. Both experiments were approved by the Departmental Ethical Committee of the University of Groningen (approval number: 16109-S-NE).

### Apparatus and Stimuli

Stimuli were controlled by Psychtoolbox, a freely available toolbox for Matlab. Visual stimuli were generated with Psychtoolbox and presented on a 17-inch (43.18 cm) CRT screen running at 100 Hz refresh rate and a resolution of 1280 by 1024 pixels. Auditory stimuli were generated with the freely available software Audacity and were presented with stereo Logitech computer speakers. The intensity of all tones was adjusted to 70 dB SPL at a fixed distance of 60 cm between speakers and participants in both experiments. All tones had 10 ms ramp up and ramp down time. Responses were collected with a custom two-button response box, connected via a USB interface.

The memory items used in the auditory WM experiment were 8 pure tones, ranging from 270 Hz to 3055 Hz in steps of half an octave. The probes in the auditory experiment were 16 pure tones that were one-third of an octave higher or lower than the corresponding auditory memory items.

The memory items used in the visual WM experiment were 8 sine-wave gratings with orientations of 11.25° to 168.75° in steps of 22.5°. The visual probes were 16 sine-wave gratings that were rotated 20° clockwise or counter-clockwise relative to the corresponding visual memory items. All gratings were presented at 20% contrast, with a diameter of 6.5° (at 60 cm distance) and a spatial frequency of 1 cycle per degree. The phase of each grating was randomized within and across trials.

The remaining stimuli were the same in both experiments. The retro-cue was a number (“1” or “2”) that subtended 0.7°. The visual impulse stimulus was a white circle with a diameter of 12°. The auditory impulse was a complex tone consisting of the combination of all pure tones used as memory items in the auditory task. A grey background (RGB = 128, 128, 128) and a black fixation dot with a white outline (0.25°) were maintained throughout the trials. All visual stimuli were presented in the centre of the screen.

### Procedure

The trial structure was the same in both experiments, as shown in Figure 1 (panels A and C). In both cases, participants completed a retro-cue WM task. Only the memory items and probes differed between experiments. Memory items and probes were pure tones in the auditory WM task and sine-wave gratings in the visual WM task. Each trial began with the presentation of a fixation dot, which stayed on the screen throughout the trial. After 1,000 ms the first memory item was presented for 200 ms. After a 700 ms delay the second memory item in the same modality as the first item was presented for 200 ms. Each memory item was selected randomly without replacement from a uniform distribution of 8 different tonal frequencies or grating orientations (see above) for the auditory and visual experiment, respectively. After another delay of 700 ms, the retro-cue was presented for 200 ms, indicating to participants whether the first or second memory item would be tested at the end of the trial. After a delay of 1,000 ms the impulse stimuli (the visual circle and the complex tone) were presented serially for 100 ms each with a delay of 900 ms in-between. The order of the impulses was fixed for each participant but counter-balanced between participants. The probe stimulus followed 900 ms after the second impulse offset and was presented for 200 ms. In the auditory WM experiment the probe was a pure tone and the participant’s task was to indicate via button press on the response box whether the probe’s frequency was lower (left button) or higher (right button) than the cued memory item. In the visual task, the probe was another visual grating, and the participants indicated whether it was rotated counterclockwise (left button) or clockwise (right button) relative to the cued memory item. The direction of the tone or tilt was selected randomly without replacement from a uniform distribution. After each response, a smiley face was shown for 200 ms, which indicated whether the response was correct or incorrect. The next trial began automatically after a randomized, variable delay of 700-1,000 ms after response input. Each experiment consisted of 768 trials in total and lasted approximately 2 hours.

**Figure 1.**
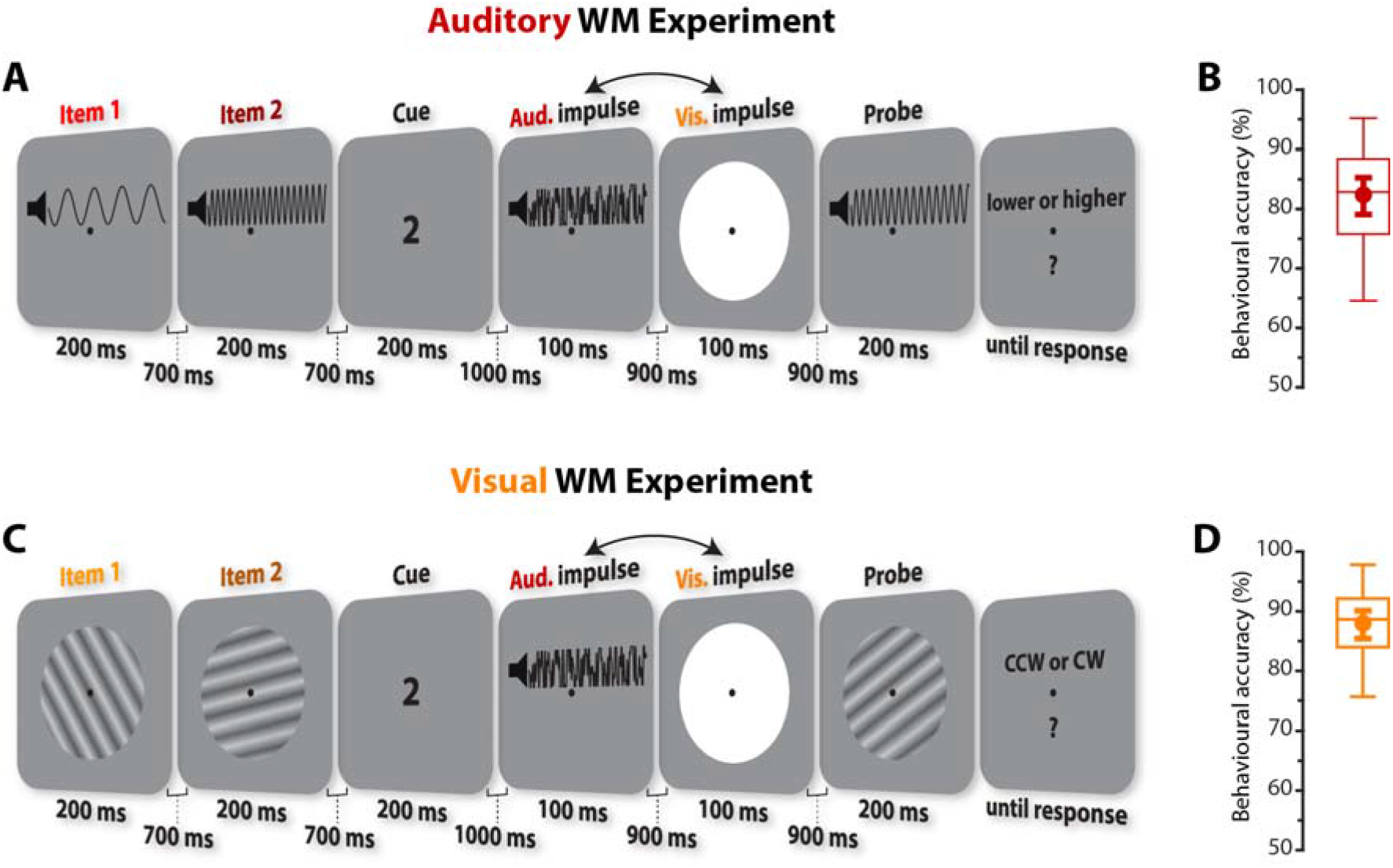
Task structure and behavioural performance. (**A**) Trial schematic of auditory task. Two randomly selected pure tones (270 Hz to 3055 Hz) were serially presented and a retro-cue indicated which of those tones would be tested at the end of the trial. In the subsequent delay, two irrelevant impulse stimuli (a complex tone and a white circle) were serially presented. At the end of each trial another pure tone was presented (the probe), and participants were instructed to indicate whether the frequency of the previously cued tone was higher or lower than the probe’s frequency. (**B**) The boxplot shows auditory task accuracy. Centre line indicates the median; box outlines show 25^th^ and 75^th^ percentiles, and whiskers indicate 1.5x the interquartile range. The superimposed circle and error bars indicate the mean and its 95% C.I., respectively. (**C**) Trial schematic of visual task. The trial structure was the same as in the auditory task. Instead of pure tones, memory items were randomly orientated gratings. The probe was another orientation grating, and participants were instructed to indicate whether the cued item’s orientation was rotated clockwise or counter-clockwise relative to the probe’s orientation. (**D**) Visual task performance.

### EEG acquisition and pre-processing

The EEG signal was acquired from 62 Ag/AgCls sintered electrodes laid out according to the extended international 10-20 system. An analog-to-digital TMSI Refa 8-64/72 amplifier and Brainvision recorder software were used to record the data at 1000 Hz using an online average reference. An electrode placed just above the sternum was used as the ground. Bipolar electrooculography (EOG) was recorded by electrodes placed above and below the right eye, and to the left and right of the left and right eye, respectively. The impedances of all electrodes were kept below 10 kΩ.

Offline the data was down-sampled to 500 Hz and bandpass filtered (0.1 Hz high-pass and 40 Hz low-pass) using EEGLAB (Delorme & Makeig, 2004). The data was epoched relative to the onsets of the memory items (-150 ms to 900 ms) and to the onsets of the auditory and visual impulse stimuli (-150 to 500 ms). The signal’s variance across channels and trials was visually inspected using a visualization tool provided by the Matlab extension Fieldtrip (Oostenveld et al., 2010), and especially noisy channels were removed and replaced through spherical interpolation, while noisy epochs were removed from all subsequent electrophysiological analyses. Epochs containing any artefacts related to eye movements were identified by visually inspecting the EOG signals and also removed from analyses.

### Multivariate pattern analysis of neural dynamics

We wanted to test if the electrophysiological activity evoked by the memory-stimuli and impulse-stimuli contained item-specific information. Since event-related potentials are highly dynamic, we used an approach that is sensitive to such changing neural activity within predefined time-windows, by pooling relative voltage fluctuations over space (i.e., electrodes) and time. This approach has two key benefits: First, pooling information over time (in addition to space) multivariately can boost decoding accuracy (Grootswagers, Wardle, & Carlson, 2017; Nemrodov, Niemeier, Patel, & Nestor, 2018). Secondly, by removing the mean-activity level within each time-window, the voltage fluctuations are normalized. This is similar to taking a neutral pre-stimulus baseline common in ERP analysis. Notably, this also removes stable activity traces that do not change within the chosen time-window, making this approach ideal to decode transient, stimulus-evoked activation patterns, while disregarding more stationary neural processes. The following details of the analyses were the same for each experiment, unless explicitly stated.

For the time-course analysis, we used a sliding window approach that takes into account the relative voltage changes within a 100 ms window. The time-points within 100 ms of each channel and trial were first down-sampled by taking the average every 10 ms, resulting in 10 voltage values for each channel. Next, the mean activity within that time-window of each channel was subtracted from each individual voltage value. All 10 voltage values per channel were then used as features for the 8-fold cross-validation decoding approach.

We used Mahalanobis distance (De Maesschalck, Jouan-Rimbaud, & Massart, 2000) to quantify the potentially parametric neural activity underlying the processing and maintenance of orientations and tones. The distances between each of the left-out test-trials and the averaged, condition-specific patterns of the train-trials (tones and orientations in the auditory and visual experiment, respectively), were computed, with the covariance matrix estimated from the train-trials using a shrinkage estimator (Ledoit & Wolf, 2004). To acquire reliable distance estimates, this process was repeated 50 times, where the data was randomly partitioned into 8 folds using stratified sampling each time. The number of trials of each condition (orientation/tone frequency) of the 7 train-folds were equalized by randomly subsampling the minimum number of condition-specific trials to ensure an unbiased training set. The average was then taken of these repetitions. For each trial the 8 distances (one of each stimulus condition) were sign-reversed for interpretation purposes, so that higher values reflect higher pattern-similarity between test and train-trials. For visualization, the sign-reversed distances were furthermore mean-centred by subtracting the mean distance of all distances of a given trial and ordered as a function of tone difference, in 1 octave steps by averaging over adjacent half octave differences, and orientation difference.

To summarize the expected positive relationship between tone-similarity and neural activation similarity (indicative of tone-specific information in the recorded signal) into a single value in the auditory WM experiment, the absolute tonal differences were linearly regressed against the corresponding pattern similarity values for each trials. The obtained beta values of the slopes were then averaged across all trials to represent “decoding accuracy”, where high values suggest a strong positive effect of tone similarity on neural pattern similarity.

To summarize the tuning curves in the visual WM experiment, we computed the cosine vector means (Wolff et al., 2017), where high values suggest evidence for orientation decoding.

The approach described above was repeated in steps of 8 ms across time (−52 to 900 ms relative to item 1 and 2 onset, and −52 to 500 ms relative to auditory and visual onset). The decoding values were averaged over trials, and the decoding time-course was smoothed with a Gaussian smoothing kernel (s.d. = 16 ms). Within the time-window, information was pooled from −100 to 0 ms relative to a specific time-point. By only including data-points from before the time-point of interest, it is ensured that decoding onsets can be more easily interpreted, whereas decoding offsets should be interpreted with caution (Grootswagers et al., 2017). In addition to the sliding window approach, we also pooled information multivariately across the whole time-window of interest (Nemrodov et al., 2018). As before, the data was first down-sampled by taking the average every 10 ms, and the mean activity from 100 to 400 ms relative to impulse onset was subtracted. The resulting 30 values per channel were then provided to the multivariate decoding approach in the same way as above, resulting in a single decoding value per participant. The time-window of interest was based on previous findings showing that the WM-dependent impulse-response is largely confined within that window (Wolff et al., 2017). Additionally, items in the item-presentation epochs were also decoded using each channel separately, using the data from 100-400 ms relative to onset. Decoding topographies were visualized using fieldtrip (Oostenveld et al., 2010).

### Cross-epoch generalization analysis

We also tested if WM-related decoding in the impulse epochs generalized to the memory presentation. Instead of using the same epoch (100-400 ms) for training and testing, as described above, the classifier was trained on the memory item epoch and tested on the impulse epoch that contained significant item decoding (and vice versa). In the auditory task, we also tested if the different impulse epochs cross-generalized by training on the visual and testing on the auditory impulse (and vice versa).

### Statistical significance testing

All statistical tests were the same between experiments. Sample sizes of all analyses were *n*=30 and *n*=28 in the auditory and visual tasks, respectively. To determine if the decoding values (see above) are higher than 0 or different between items, we used a non-parametric sign-permutation test (Maris & Oostenveld, 2007). The sign of the decoding value of each participant was randomly flipped 100.000 times with a probability of 50%. The *p* value was derived from the resulting null distribution. The above procedure was repeated for each time-point for time-series results. A cluster based permutation test (100.000 permutations) was used to correct for multiple comparisons over time using a cluster forming and cluster significance threshold of *p* < 0.05. Complementary Bayes factors to test for decoding evidence for the cued and uncued items within each impulse epoch separately were also computed.

We were also interested if there were differential effects on the decoding results between cueing (cued/uncued) and impulse modality (auditory/visual) during WM maintenance. To test this, we computed the Bayes factors of models with and without each of these predictors versus the null model that only included subjects as a predictor (Bayesian equivalent of repeated measures ANOVA). The freely available software package JASP (JASP Team, 2018) was used to compute Bayes factors.

## Results

### Parametric processing of visual and auditory stimuli

#### Auditory WM task

The neural dynamics of auditory stimulus processing showed a clear parametric effect, with a positive relationship between tone and pattern similarity (Fig. 2A & C) for both memory items. The neural dynamics showed significant item-specific decoding clusters during, and shortly after, corresponding item presentation for item 1 (44 to 708 ms relative to item 1 onset, *p* < 0.001, one-sided, corrected) and item 2 (28 to 572 ms relative to item 2 onset, *p* < 0.001, one-sided, corrected; Fig. 2B). The topographies of channel-wise item-decoding for each item using the neural data from 100-400 ms after item-onset, revealed strong decoding for frontal-central and lateral electrodes (Fig. 2D), suggesting that the tone-specific neural activity is most likely generated by the auditory cortex (Chang, Bosnyak, & Trainor, 2016). These results provide evidence that stimulus-evoked neural activity fluctuations contain information about presented tones that can be decoded from EEG.

**Figure 2.**
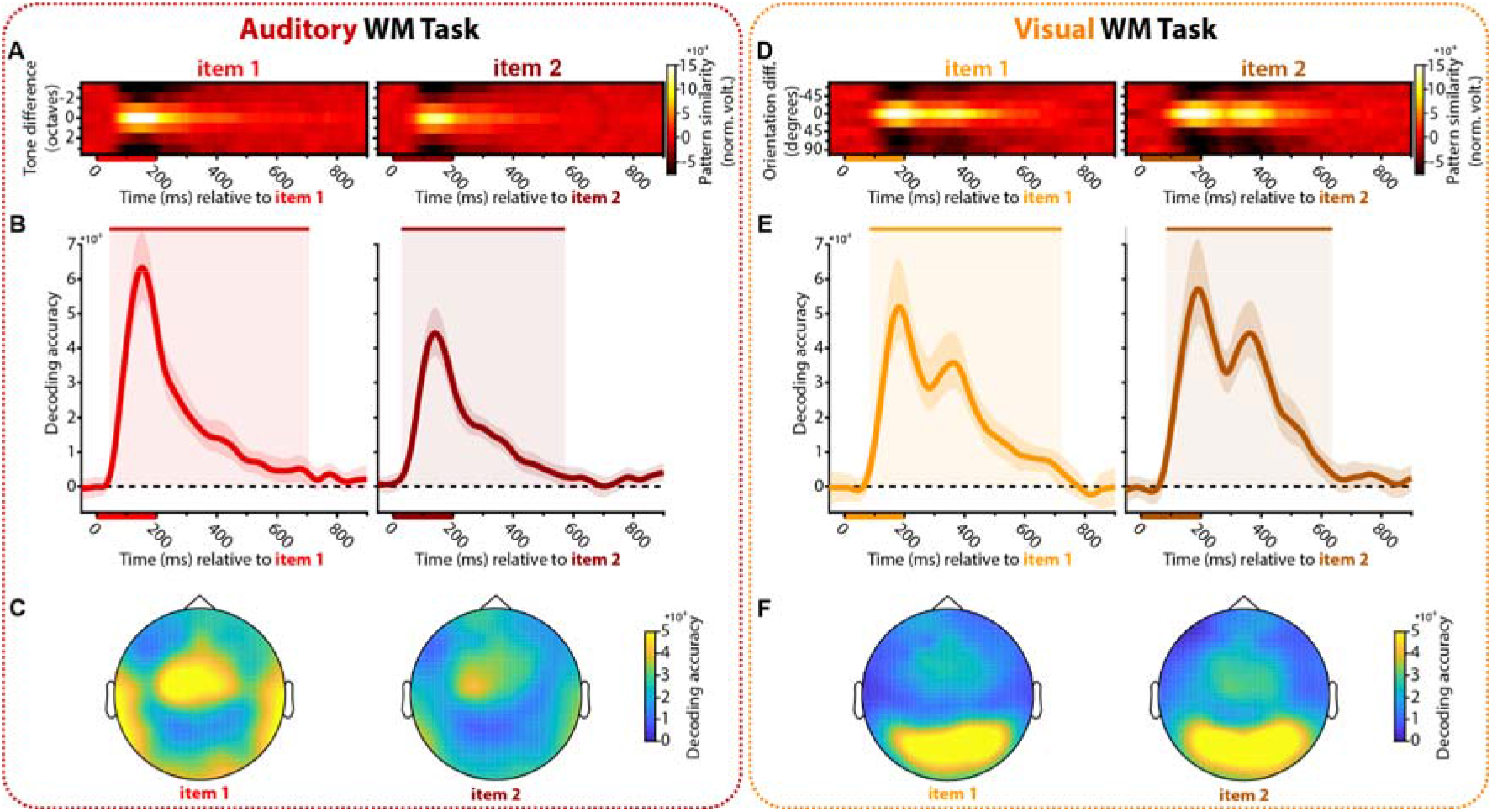
Decoding during item encoding. **A-C**, Auditory WM task. **D-F**, Visual WM task. **A & D**, Normalized average pattern similarity (mean-centred, sign-reversed mahalanobis distance) of the neural dynamics for each time-point between trials as a function of tone similarity in **A** and orientation similarity in **D**, separately for item 1 and item 2, in item 1 and item 2 epochs, respectively. Bars on the horizontal axes indicate item presentations. **B & E**, Beta values in **B** and cosine vector-means in E of pattern similarities for item 1 and 2. Upper bars and corresponding shading indicate significant values. Error shading indicates 95% C. I. of the mean. **C & F**, Topographies of each item of channel-wise decoding (100-400 ms relative to item onset).

#### Visual WM task

Processing of visual orientations also showed a parametric effect (Fig. 2E & G), replicating previous findings (Saproo & Serences, 2010). The item-specific decoding time-courses of the dynamic activity showed significant decoding clusters during and shortly after item presentations (item 1: 84-724 ms, *p* < 0.001; item 2: 84-636 ms, *p* < 0.001, one-sided, corrected). As expected, the topographies of channel-wise item-decoding showed strong effects in posterior channels that are associated with the visual cortex.

### Content-specific impulse responses

#### Auditory WM task

In the auditory impulse epoch, the neural dynamics time-course revealed significant cued-item decoding (180-252 ms, *p* = 0.005, one-sided, corrected), while no clusters were present for the uncued item (Fig. 3A & B, left). Similarly, the cued item was decodable in the visual impulse epoch (204-372ms, *p* = 0.009, one-sided, corrected), while the uncued item was not (Fig. 3A & B, right).

**Figure 3.**
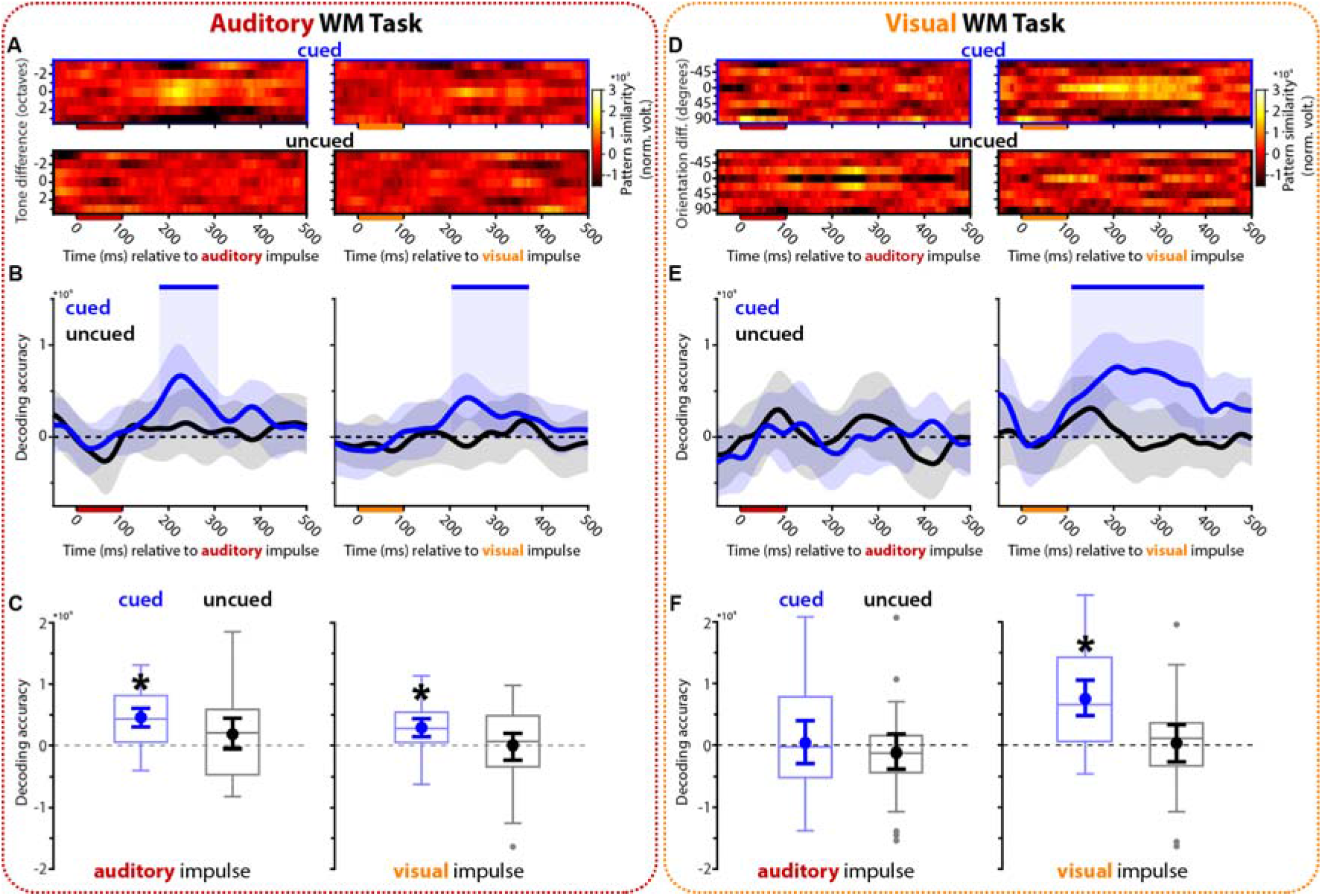
Decoding auditory and visual WM content from the impulse response. **A-C**, Auditory WM task. **D-F**, Visual WM task. **A & D**, Normalized average pattern similarity (mean-centred, sign-reversed mahalanobis distance) of the neural dynamics for each time-point between trials as a function of tone similarity in **A** and orientation similarity in **D**. Top row: cued item. Bottom row: uncued item. Left column: auditory impulse. Right column: visual impulse. **B & E**, Decoding accuracy time-course: Beta values in **B** and cosine vector-means in **E** of pattern similarities for cued (blue) and uncued item (black). Upper bars and shading indicate significant values of the corresponding item. Error shading indicate 95% C. I. of the mean. **C & F**, Boxplots show the overall decoding accuracies for the cued (blue) and uncued (black) item, using the whole time-window of interest (100-400 ms relative to onset) from the auditory (left) and visual (right) impulse epoch. Centre lines indicate the median; box outlines show 25^th^ and 75^th^ percentiles, and whiskers indicate 1.5x the interquartile range. Extreme values are shown separately (dots). Superimposed circles and error bars indicate mean and its 95% C.I., respectively. Asterisks indicate significant decoding accuracies (*p* < 0.05, one-sided).

The time-of-interest (100-400 ms relative to impulse onset) analysis provided similar results. The cued item showed strong decoding in both impulse epochs (auditory impulse: *p* < 0.001, Bayes factor = 11462.607; visual impulse: *p* < 0.001, Bayes factor = 85.843, one-sided), but the uncued item did not (auditory impulse: *p* = 0.075, Bayes factor = 0.968; visual impulse: *p* = 0.476, Bayes factor = 0.204, one-sided; Fig. 3C). A model only including the cueing predictor yielded the highest Bayes factor of 8.120 (± 2.104 %) compared to the null model. A model including impulse modality as a predictor resulted in a Bayes factor of 0.829 (± 1.255 %). Including both predictors (impulse modality and cueing) in the model resulted in a Bayes factor of 7.511 (± 2.213 %) that was slightly lower than only including cueing.

Taken together, these results provided strong evidence that both impulse stimuli elicit neural responses that contain information about the cued item in auditory WM, but none about the uncued item.

#### Visual WM task

No significant time clusters were present in the auditory impulse epoch of the visual WM experiment for either the cued or the uncued item task (Fig. 3D & E, left). The decoding time-course of the visual impulse epoch revealed a significant decoding cluster of the cued item (108-396 ms, *p* <0.001, one-sided, corrected) but not for the uncued item (Fig. 3D & E, right), replicating previous findings (Wolff et al., 2017).

The analysis on the time-of-interest interval (100-400 ms) showed the same pattern of results; neither the cued, nor uncued item in the auditory impulse epoch showed above 0 decoding (cued: *p* = 0.417, Bayes factor = 0.236; uncued: *p* = 0.787, Bayes factor = 0.119, one-sided). In the visual impulse epoch the cued item showed strong decodability (*p* < 0.001, Bayes factor = 1695.823, one-sided) but the uncued item did not (*p* = 0.421, Bayes factor = 0.119, one-sided; Fig 3F). A model including both predictors (cueing and impulse modality) as well as their interaction resulted in the highest Bayes factor compared to the null model (Bayes factor = 55.728 (± 2.155 %)). Models with each predictor alone resulted in notably smaller Bayes factors (cueing: Bayes factor = 6.191 (± 0.829 %); impulse modality: Bayes factor = 5.853 (± 1.116 %)). The Bayes factor of the model including both predictors without interaction (47.032 (± 2.155 %)) was only 1.185 times smaller than the model that also included the interaction, highlighting that while there was strong evidence in favour of both impulse modality and cueing, there was only weak evidence in favour of an interaction.

Overall, these results provided evidence that while a visual impulse clearly evokes a neural response that contains information about the cued visual WM item, replicating previous findings (Wolff et al., 2017), an auditory impulse does not.

### No WM-specifîc cross-generalization between impulse and WM-item presentation

It has been shown previously that the visual WM-dependent impulse-response does not cross-generalize with visual item processing (Wolff et al., 2015). Here we tested if this is also the case for auditory WM, and additionally explored the cross-generalizability between impulses.

#### Auditory WM task

The representation of the cued item did neither cross-generalise between item presentation and either of the impulse epochs (auditory impulse: *p* = 0.58, Bayes factor = 0.225; visual impulse: *p* = 0.26, Bayes factor = 0.356, two-sided), nor between impulse epochs (*p* = 0.417, Bayes factor = 0.267; Fig. 4A).

**Figure 4.**
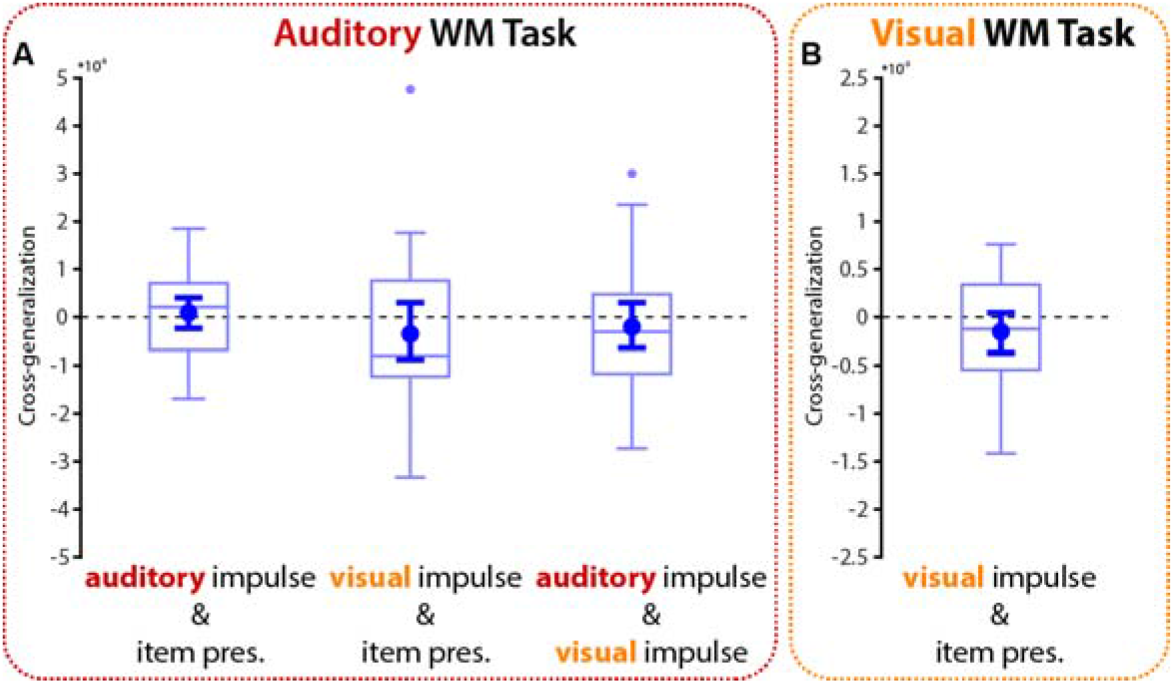
Cross-generalization between epochs. **A**, Cross-generalization of the cued item between the memory item epoch and impulse epochs in the auditory WM task. **B**, Cross-generalization between visual impulse and memory item in the visual WM task. Centre lines indicate the median; box outlines show 25^th^ and 75^th^ percentiles, and whiskers indicate 1.5x the interquartile range. Extreme values are shown separately (dots). Superimposed circles and error bars indicate mean its 95% C.I, respectively.

#### Visual WM task

Replicating previous reports (Wolff et al., 2015, 2017), the visual impulse response of the cued visual item did not cross-generalize with item processing during item presentation (*p* = 0.168, Bayes factor = 0.491).

## Discussion

It has previously been shown that the bottom-up neural response to a visual impulse presented during the delay of a visual WM task contains information about relevant visual WM content (Wolff et al., 2015, 2017), which is consistent with a key prediction of WM theories that assume information is maintained in activity-silent brain states (Stokes, 2015). We used this approach to investigate whether sensory information is maintained within sensory-specific neural networks, shielded from other sensory processing areas. We show that the neural impulse response to sensory-specific stimulation is WM content-specific not only in visual WM, but also in auditory WM, demonstrating the feasibility and generalisability of the approach in the auditory domain. Furthermore, for auditory WM, a content-specific response was obtained not only during auditory, but also during visual stimulation, suggesting a sensory modality-unspecific path to access the auditory WM network. In contrast, only visual, but not auditory, stimulation evoked a neural response containing relevant visual WM content. This pattern of impulse responsivity supports the idea that visual pathways may be more dominant in WM maintenance.

Recent studies have shown that delay activity in the auditory cortex reflects the content of auditory WM (Huang et al., 2016; Kumar et al., 2016; Uluç et al., 2018). Thus, similar to visual WM maintenance, which has been found to result in content-specific delay-activity in the visual cortex (Harrison & Tong, 2009), auditory WM content is also maintained in a network that recruits the same brain area responsible for sensory processing. However, numerous visual WM studies have shown that content-specific delay activity may in fact reflect the focus of attention (Lewis-Peacock, Drysdale, Oberauer, & Postle, 2011; Sprague, Ester, & Serences, 2016; Watanabe & Funahashi, 2014). The memoranda themselves may instead be represented within connectivity patterns that generate a distinct neural response profile to internal or external neural stimulation (Lundqvist et al., 2016; Rose et al., 2016; Wolff et al., 2017). While previous research has focused on visual WM, we now provide evidence for a neural impulse response that reflects parametric auditory WM content, suggesting a similar neural mechanism for auditory WM.

The neural response to a visual impulse stimulus also contained information about the behaviourally relevant pitch. It has been shown that visual stimulation can result in neural activity in the auditory cortex (Martuzzi et al., 2007; Morrill & Hasenstaub, 2018). Thus, direct connectivity between visual and auditory areas (Eckert et al., 2008) might be such that visual stimulation activates auditory WM representations in auditory cortex, providing an alternate access pathway. Alternatively, visual cortex itself might retain auditory information. It has previously been shown that natural sounds can be decoded from the activity in the visual cortex, during both processing and imagination (Vetter, Smith, & Muckli, 2014). Even though pure tones were used in the present study, it is nevertheless possible that they have been visualised, for example by imagining the pitch as a location in space. Tones may have also resulted in semantic representations, by categorising them into arbitrary sets of low, medium, and high tones. The decodable signal from the impulse-response might thus not necessarily originate from the sensory-processing areas, but rather from higher brain regions such as the prefrontal cortex (Stokes et al., 2013). Future studies that employ imaging tools with high spatial resolution might be able to arbitrate the neural origin of the cross-modal impulse response in WM.

While the neural impulse response to visual stimulus contained information about the relevant visual WM item, replicating previous results (Wolff et al., 2017), the neural response to external auditory stimulation did not. This suggests that, in contrast to auditory information, visual information is maintained in a sensory-specific neural network with no evidence of content-specific connectivity with the auditory system, possibly reflecting the visual dominance of the human brain (Posner, Nissen, & Klein, 1976). Indeed, while it has been found that auditory stimulation results in neural activity in the visual cortex, it is notably weaker than the other way around (Martuzzi et al., 2007), which corresponds with our asymmetric findings of sensory specific and sensory non-specific impulse responses of visual and auditory WM between visual and auditory cortices.

It has previously been reported that the WM-related neural pattern evoked by the impulse response does not cross-generalize with the neural activity evoked by the memory stimulus itself (Wolff et al., 2015), suggesting that the neural activation patterns are qualitatively different. In the present study, we also found no cross-generalization between item processing and the impulse response, neither in the visual nor in the auditory WM task. The neural representation of WM content may thus not be an exact copy of stimulation history, literally reflecting the activity pattern during information processing and encoding, but rather a reconfigured code that is optimized for future behavioural demands (Myers, Stokes, & Nobre, 2017). Similarly, no generalizability was found between auditory and visual impulse responses in the auditory task. This could suggest that distinct neural networks are perturbed by the different impulse modalities, or, as alluded to above, that it reflects the unique interaction between impulses and the perturbed neural network. Future research should employ neural imaging tools with high spatial resolution to investigate the neural populations involved in the WM-dependent impulse-response.

The present results provide a novel approach to the ongoing debate on the extent to which sensory processing areas are essential for the maintenance of information in WM (Gayet, Paffen, & Van der Stigchel, 2018; Scimeca et al., 2018; Xu, 2018). This is usually investigated by looking for the presence of WM-specific delay activity in the visual cortex in visual WM tasks (Bettencourt & Xu, 2016; Harrison & Tong, 2009), where null-results are interpreted as evidence against the involvement of specific brain regions, which is inherently problematic (Ester, Rademaker, & Sprague, 2016), and by which non-active WM states are not considered. In the present study, we found that sensory-specific stimulation, and both sensory specific and non-specific stimulation, resulted in WM-specific neural responses during the maintenance of visual and auditory information, respectively. Sensory cortices were thus linked to WM maintenance not by relying on ambient delay-activity, but rather by perturbing the underlying, connectivity-dependent, representational WM network via a bottom-up neural response.

## Acknowledgments

This research was in part funded by a Biotechnology and Biological Sciences Research Council (BB/M010732/1) and James S. McDonnell Foundation Scholar Award (220020405) to MGS, and by the NIHR Oxford Health Biomedical Research Centre. The Wellcome Centre for Integrative Neuroimaging is supported by core funding from the Wellcome Trust (203139/Z/16/Z). The views expressed are those of the authors and not necessarily those of the National Health Service, the National Institute for Health Research or the Department of Health. We would like to thank P. Albronda for providing technical support and M. Rietdijk for helping with data collection.

